# Strengthening E-cadherin adhesion via antibody mediated binding stabilization

**DOI:** 10.1101/2023.07.04.547716

**Authors:** Bin Xie, Shipeng Xu, Leslayann Schecterson, Barry M. Gumbiner, Sanjeevi Sivasankar

**Affiliations:** Biophysics Graduate Group, University of California, Davis, CA; Department of Biomedical Engineering, University of California, Davis, CA; Seattle Children’s Research Institute, Center for Developmental Biology and Regenerative Medicine, Seattle, WA.; Department of Cell Biology, University of Virginia, Charlottesville, VA.

## Abstract

E-cadherins (Ecads) are a crucial cell-cell adhesion protein with tumor suppression properties. Ecad adhesion can be enhanced by the monoclonal antibody 66E8, which has potential applications in inhibiting cancer metastasis. However, the biophysical mechanisms underlying 66E8 mediated adhesion strengthening are unknown. Here, we use molecular dynamics simulations, site directed mutagenesis and single molecule atomic force microscopy experiments to demonstrate that 66E8 strengthens Ecad binding by stabilizing the primary Ecad adhesive conformation: the strand-swap dimer. By forming electrostatic interactions with Ecad, 66E8 stabilizes the swapped β-strand and its hydrophobic pocket and impedes Ecad conformational changes, which are necessary for rupture of the strand-swap dimer. Our findings identify fundamental mechanistic principles for strengthening of Ecad binding using monoclonal antibodies.

## Introduction

E-cadherins (Ecads) are essential cell-cell adhesion proteins that mediate epithelial tissue formation, regeneration, and differentiation (Honig and Shapiro, 2020). Deficient Ecad adhesion leads to a loss of contact inhibition and increased cell mobility, contributing to the metastasis of various types of cancer (Mendonsa et al., 2018; Petrova et al., 2016).

Ecad adhesion is mediated by calcium dependent, *trans* homophilic interactions of opposing Ecad extracellular regions (ectodomains). Ecad’s primary adhesive conformation, called a strand-swap dimer, is formed by the exchange of N-terminal β-strands between the outermost domains (EC1) of opposing Ecads. This exchange of β-strands results in the symmetric docking of a conserved W2 anchor residue into a complementary pocket on the partner Ecad (Brasch et al., 2012; Haussinger et al., 2004; Parisini et al., 2007). Stabilizing β-strands (Vendome et al., 2011) and their complementary binding pockets (Vendome et al., 2014b) thermodynamically stabilizes the strand-swap dimer. However, the molecular details of how the stability of the β-strand and pocket can be extrinsically controlled and why this process strengthens adhesion are unclear.

Strengthening Ecad adhesion is not just interesting at the fundamental biophysical level but also has potential therapeutic significance. Due to Ecad’s role as a tumor suppressor, there is ongoing interest in developing strategies to activate or reinforce Ecad adhesion, and thereby mitigate cancer metastasis. One promising therapeutic approach is the development of monoclonal antibodies (mAbs) that can modulate the interaction of Ecad. Adhesion activating mAbs, such as 66E8 and 19A11, have been developed to specifically target the Ecad ectodomains and reinforce cell-cell adhesion (Petrova et al., 2012). In mouse models, these mAbs have been shown to effectively impede the metastatic invasion of lung cancer cells expressing human Ecad (Na et al., 2020). Similar mAb based approaches have also been deployed to specifically target other adhesion proteins such as integrins for the treatment of Crohn’s disease, (Byron et al., 2009; Ley et al., 2016; Raab-Westphal et al., 2017).

We recently showed that 19A11 strengthens Ecad adhesion by forming salt bridges that stabilize the β-strand and its complementary binding pocket (Xie et al., 2022). Similarly, cryo-EM suggests that Ecad bound to 19A11 adopts a twisted conformation which may represent a strengthened strand-swap dimer (Maker et al., 2022). In contrast to 19A11 which binds to the base of the swapped β-strand on the EC1 domain, a recent crystal structure shows that 66E8 binds to the Ecad EC2 domain and exhibits no interactions with the Ecad EC1 domain (Maker et al., 2022). However, surprisingly, 66E8 still influences strand-swap dimer formation, which involves the N-terminal EC1 domain. This raises questions on whether 66E8 stabilizes the Ecad β-strands and hydrophobic pockets and how this ultimately strengthens Ecad binding. Addressing these questions will establish guidelines for rationally designing mAbs that strengthen Ecad mediated cell adhesion.

Here, we determine the molecular mechanism underlying 66E8 mediated strengthening of Ecad binding. Using atomistic computer simulations and single molecule atomic force microscopy (AFM) experiments, we show that 66E8 forms a novel interface with the Ecad EC1 domain, an interface that is not present in the crystal structure. Establishment of this *de novo* interface results in formation of a salt bridge and hydrogen bonds between 66E8 and the base of each swapped β-strand in Ecad, which stabilizes the β-strand and the hydrophobic pocket. Importantly, these interactions between 66E8 and Ecad impede conformational changes, which are necessary for rupture of the strand-swap dimer. Our findings identify fundamental mechanistic principles for strengthening of Ecad binding using mAbs.

## Results

### 66E8 forms novel interface with Ecad EC1 domain which stabilizes the strand-swap dimer

The crystal structure of 66E8 Fab binding on Ecad has been solved recently (PDB ID code 6VEL) (Maker et al., 2022). This structure shows that binding of 66E8 does not cause gross conformational changes on Ecad and the root mean square deviation (RMSD) between crystal structures in the presence and absence (PDB ID code 2O72) of 66E8 is only 0.75Å. However, upon binding to 66E8 the Ecad β-strand and the anchoring W2 residue reorients, suggesting 66E8 may affect the stability of the β-strand (Maker et al., 2022). In the crystal structure, 66E8 recognizes the Ecad EC2 domain but does not interact with Ecad EC1 domain (Figure 1a). The closest distance observed between α-carbons on the EC1 domain and 66E8 (K14:G87) is 12.2Å (Figure 1a). This suggests that the action of 66E8 is allosteric, since it binds to the Ecad EC2 domain, but affects the strand-swap dimer which involves EC1 domains.

**Fig. 1:**
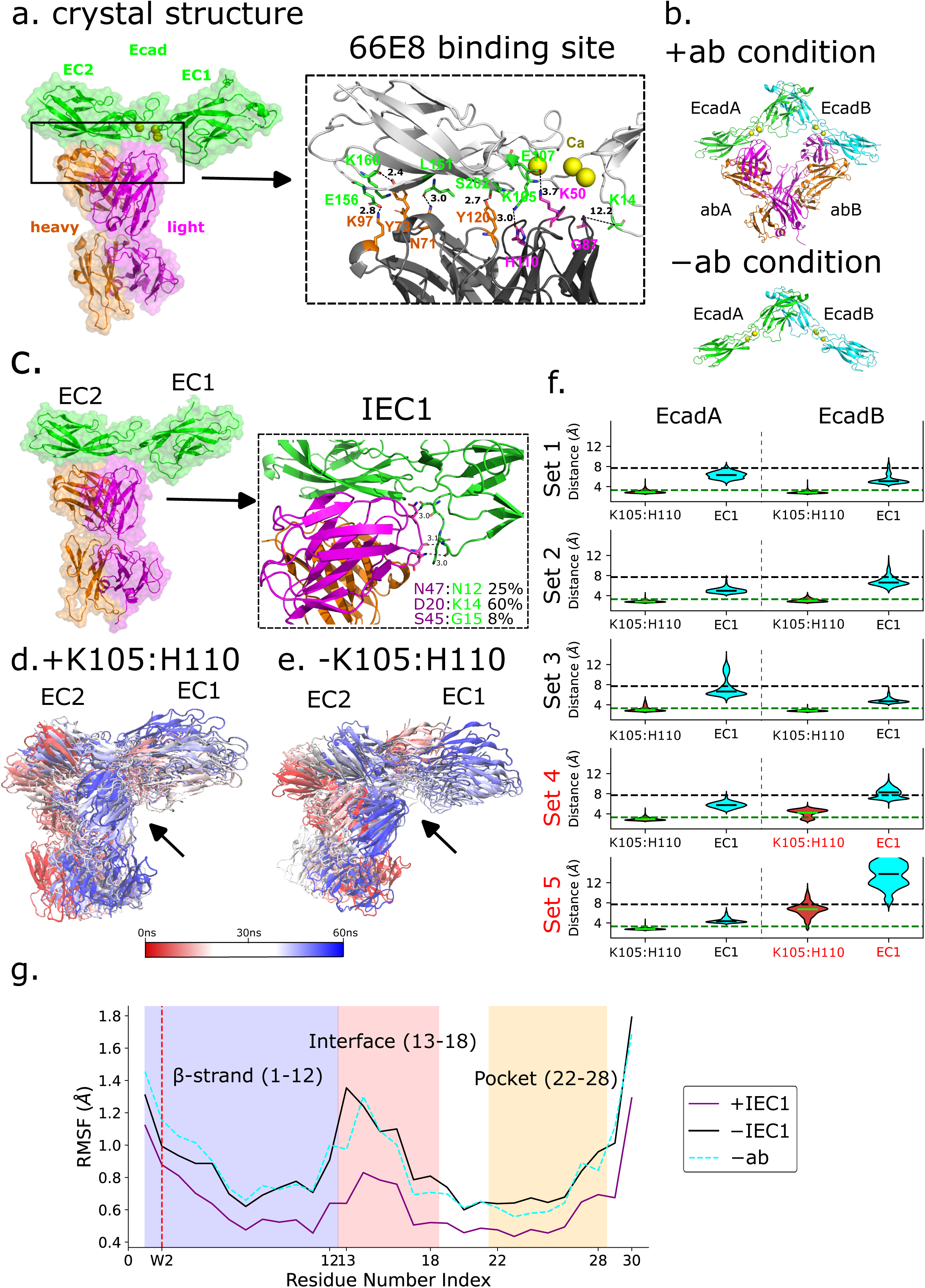
Binding of 66E8 stabilizes both the Ecad β-strand and the W2 hydrophobic pocket through the formation of interface EC1 (IEC1). (a) X-ray crystal structure of 66E8 Fab heavy chain (orange) and light chain (magenta) bound to Ecad EC1-2 domains (green). Detailed view of salt bridges or hydrogen bonds between 66E8 and Ecad EC2 domain are shown in the inset. The distance between interacting atoms are in Å (black dashed lines). (b) MD simulations were performed in two conditions: Ecad strand-swap dimer (EcadA and EcadB) in the absence of 66E8 (−ab), and Ecad strand-swap dimer with two 66E8 Fabs (abA and abB) bound to both Ecads (+ab). (c) A structure snapshot shows that 66E8 forms a novel interface with Ecad EC1 domain (IEC1) during MD simulations. Three of the most likely electrostatic interactions on this interface are shown in the inset. The electrostatic interaction pair D20:K14 accounts for 60% of all of the electrostatic interactions. (d) Hydrogen bond K105:H110 facilitates formation of IEC1 (indicated by arrow) during MD simulations. (e) IEC1 was not observed (indicated by arrow) when K105:H110 was not formed. For better visibility in *(d)* and *(e)*, only 20 frames of one simulation for each condition are displayed and frames at the start of simulation are in red, whereas frames at the end are in blue. (f) Violin plots of the distances between interacting atoms in K105:H110 hydrogen bonds and between the center of masses of opposing IEC1 interface residues measured during the last 40 ns of each +ab MD simulations. K105:H110 median distances are shown as green line while IEC1 centers of mass median distances are shown as black line on the violin plot. Distances for EcadA and EcadB are shown in the left and right panels, respectively. Distances measured for K105:H110 are shown in red, while distances measured for center of masses of IEC1 residues are shown in cyan. From top to bottom, each row represents a simulation repeats, from simulation 1 (set 1) to simulation 5 (set 5). Both EcadA and EcadB form IEC1 when hydrogen bond K105:H110 persists in sets 1-3. However, IEC1 only forms on EcadA but not on EcadB when hydrogen bond K105:H110 is not formed in set 4 and 5. (g) Comparison of the average α-carbons RMSF values when an Ecad forms IEC1 with its corresponding 66E8 (purple solid line), when an Ecad does not form IEC1 with its corresponding 66E8 (solid black line), or in the absence of 66E8 (dashed cyan line). The W2 position is highlighted using a vertical dashed red line. The lower RMSF values show the formation of interface EC1 between 66E8 and Ecad stabilizes the Ecad β-strand and the hydrophobic pocket.

To understand the molecular basis by which 66E8 upregulates Ecad binding, we performed molecular dynamics (MD) simulations using two conditions: Ecad strand-swap dimer (EcadA and EcadB) without 66E8 Fab (Figure 1b bottom, “−ab” condition, PDB ID code 2O72), and Ecad strand-swap dimer bound to two 66E8 Fabs (Figure 1b top, “+ab” condition, PDB ID code 6VEL). For each condition we performed 5 repeat MD simulations; each repeat simulation was run for 60ns until the system equilibrated (Supplementary Fig. S1). Our MD simulations showed that a novel interface, henceforth called ‘interface EC1’ or ‘IEC1’, that is different from the 66E8 binding interface, was formed between 66E8 light chain and Ecad EC1 domain (Figure 1c). IEC1 was formed by 66E8 light chain residues 20 and 45-47 interacting with Ecad EC1 loop residues 13-18, which lie in between the β-strands (residues 1-12) and partial hydrophobic pocket (residues 22-28). This suggests that the formation of IEC1 could affect the Ecad strand-swap binding conformation. We characterized all salt bridges and hydrogen bonds that mediate IEC1 and found that the residue pair D20:K14 was the predominant interaction, accounting for 60% of all electrostatic interactions observed on IEC1 (Figure 1c). This finding suggests that the formation and stability of IEC1 is highly dependent on the Ecad K14 residue.

The formation of IEC1 relies on a crucial hydrogen bond between K105 and H110 observed in the crystal structure (Figure 1a), which lies adjacent to the Ecad EC1 domain (hydrogen bonds were assumed to form when the distance between interacting atoms was below 3.3Å). Formation of the K105:H110 hydrogen bond closed the gap between 66E8 and Ecad EC1 and caused 66E8 to move towards the EC1 domain and form IEC1 (Figure 1d, indicated by an arrowhead). Conversely, when the K105:H110 hydrogen bond was not formed, 66E8 remained distant from the Ecad EC1 domain (Figure 1e, indicated by an arrowhead). To assess the formation of IEC1, we monitored the distance between the centers of mass of interface residues throughout the MD simulations assuming that IEC1 was formed when the median distance between the centers of mass was below 7.5Å. (Salamanca Viloria et al., 2017). Our data showed that in 60% of the +ab simulations (sets 1-3, Fig. 1f), both Ecads form IEC1 with the bound 66E8. However, in the remaining 40% of the simulations, only one of the Ecad formed an IEC1 with 66E8, and the other Ecad did not form IEC1 due to the absence of hydrogen bonds K105:H110 (EcadB of set 4, EcadB of set 5, highlighted in red, Fig. 1f).

To evaluate the impact of IEC1 on the stability of Ecad’s β-strand and its complementary binding pocket, we measured the root-mean-square fluctuations (RMSF) of the corresponding α-carbon residues during the final 35 ns of all MD simulations. A comparison of the average RMSF values between cases where IEC1 was formed and cases where it was not formed revealed notable differences. When IEC1 was formed (Figure 1g, purple line), the RMSF of the interacting loop (residues 13-18), the β-strand and the complementary pocket decreased indicating that these motifs were stabilized. Conversely, when IEC1 was not formed (Figure 1g, black line), the stability of the protein remained similar to the -ab condition (Figure 1g, dashed cyan line). Based on these findings, we concluded that 66E8 can interact with Ecad in two distinct modes: one involving the formation of IEC1 which stabilizes the β-strand and the pocket region, and the other where IEC1 is not formed which leaves the protein stability unaffected.

### Ecad K14E mutants diminish the 66E8 stabilizing effect

To confirm the role of IEC1 in stabilizing the strand-swap dimer, we proceeded to disrupt IEC1 while leaving the 66E8 binding interface unaffected. Since the electrostatic interaction D20:K14 was the most prominent interaction observed on IEC1, we replaced the positively charged K14 on Ecad with a negatively charged glutamic acid. To evaluate the impact of the K14E mutation, we conducted MD simulations (5 repeats) of 66E8 bound to the Ecad K14E mutants. While 66E8 still formed an interface with the K14E mutant EC1 domain (Figure 2a, denoted as IEC’), the mutation eliminated the D20-K14 interaction. No new electrostatic interactions were introduced at this interface as a result of the K14E mutation (Figure 2a).

**Figure 2.**
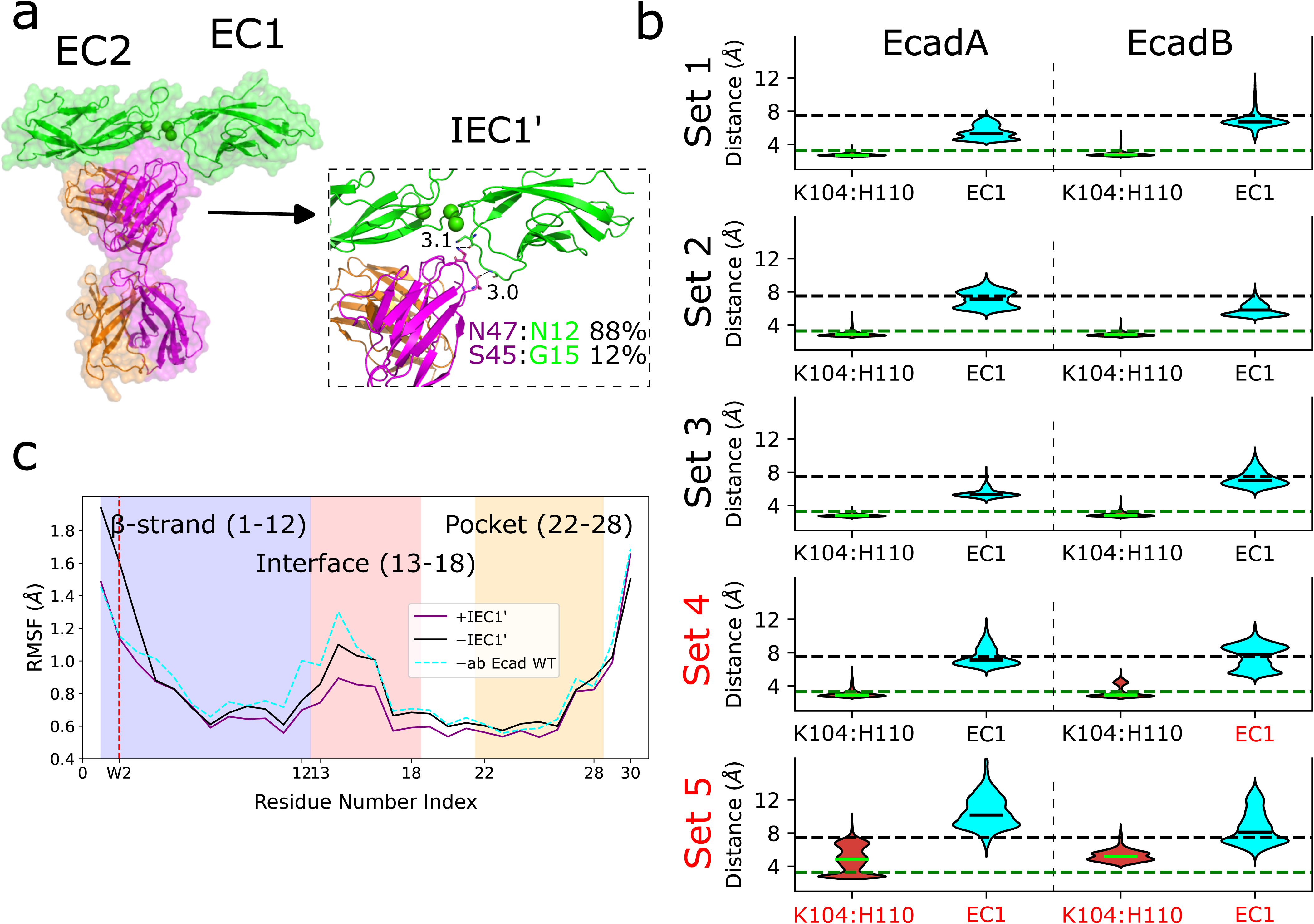
**Ecad K14E mutants decreases 66E8 induced stabilization of Ecad β-strand and hydrophobic pocket**. (a) Snapshot of 66E8 bound to Ecad K14E mutants during MD simulations. An interface between 66E8 and the K14E mutant still forms (interface EC1’ or IEC’). Detailed view of electrostatic interactions observed on IEC1’ is shown inset. Only two hydrogen bonds are observed on this interface, and K14E mutation does not introduce additional electrostatic interactions. (b) Violin plots of the distances between interacting atoms in K105:H110 hydrogen bonds and center of masses of opposing interface IEC1’ residues measured during the last 40 ns of each +ab MD simulations. Median K105:H110 distances are shown as green line while median IEC’ distances shown as black line on the violin plot. Distances for EcadA and EcadB are shown in the left and right panels, respectively. Distances measured for K105:H110 are shown in red, and distance between center of mass of IEC1’ residues are shown in cyan. From top to bottom, each row represents a simulation repeat, from simulation 1 (set 1) to simulation 5 (set 5). Both EcadA and EcadB form IEC1’ when hydrogen bond K105:H110 persists in sets 1-3. However, IEC1’ only forms on EcadA but not on EcadB even though hydrogen bond K105:H110 forms on both Ecads in set 4. In set 5, IEC1’ is not formed on both Ecads when hydrogen bond K105:H110 is not formed on both sides. (c) Comparison of the average α-carbons RMSF values when the K14E mutant forms IEC1’ with its corresponding 66E8 (purple solid line), when the K14E mutant does not forms IEC1’ with its corresponding 66E8 (solid black line), and for WT Ecad in the absence of 66E8 (dashed cyan line). The W2 position is highlighted using a vertical dashed red line. The formation of IEC1’ slightly lowers the RMSF values due to partial stabilization of β-strand and hydrophobic pocket.

We also measured the formation of IEC1’ by measuring the distances between interface residues’ center of mass and by recording the formation of K105:H110 hydrogen bond. In three simulations (set 1, 2 and 3), formation of the K105:H110 hydrogen bond triggered establishment of IEC1’ (Figure 2b, black sets). However, in sets 4 and 5, at least one of the Ecads failed to form a stable IEC1’. Specifically, in set 4, EcadB did not form IEC1’ despite the presence of the hydrogen bond K105:H110. Additionally, in set 5, neither EcadA nor EcadB stably formed IEC1’ due to the absence of stable K105:H110 hydrogen bonds on both Ecads. These findings suggest that the K14E mutants destabilize the interface formation between 66E8 and Ecad EC1 domain.

To assess the impact of the K14E mutation on 66E8 mediated stabilization of the β-strand and its complementary pocket, we analyzed the RMSF of the corresponding α-carbon residues during the final 35 ns of all MD simulations. Comparing the average RMSF values between cases where IEC1’ was formed and cases where it was not formed, we observed less stabilization compared to WT-Ecad (Figure 2C) suggesting that the K14E mutation decreases the stabilization on the β-strand and complementary pocket.

### 66E8 mediated stabilization of strand-swap dimer leads to stronger Ecad binding

To test if stabilization of strand-swap dimers by 66E8 strengthens Ecad binding, we conducted SMD simulations on structures obtained from the final frame of every MD simulation. In each SMD simulation, we immobilized the C-terminal of an Ecad molecule and applied a constant force of ∼665 pN to the C-terminus of the other Ecad molecule (Supplementary movie 1). During the SMD simulations, we monitored the interfacial binding area between the two Ecads by measuring the change in solvent accessible surface area (ΔSASA) (Marsh and Teichmann, 2011), where a decrease in ΔSASA to zero indicated the rupture of the interacting *trans* dimer. In the -ab condition (Fig. 3a), the ΔSASA reached zero at ∼1000ps, indicating the dissociation of the interacting *trans* dimer. In contrast, the +ab condition exhibited two distinct populations: one population remained bound for a longer duration, while the other population unbound within a similar timescale as the -ab condition. Notably, in sets 1, 2, and 3 of the +ab condition, the Ecads interacted for ∼3000ps (Fig. 3b), indicating a robust bound state. However, in sets 4 and 5, the interactions lasted only ∼1200ps (Fig. 3b).

**Figure 3.**
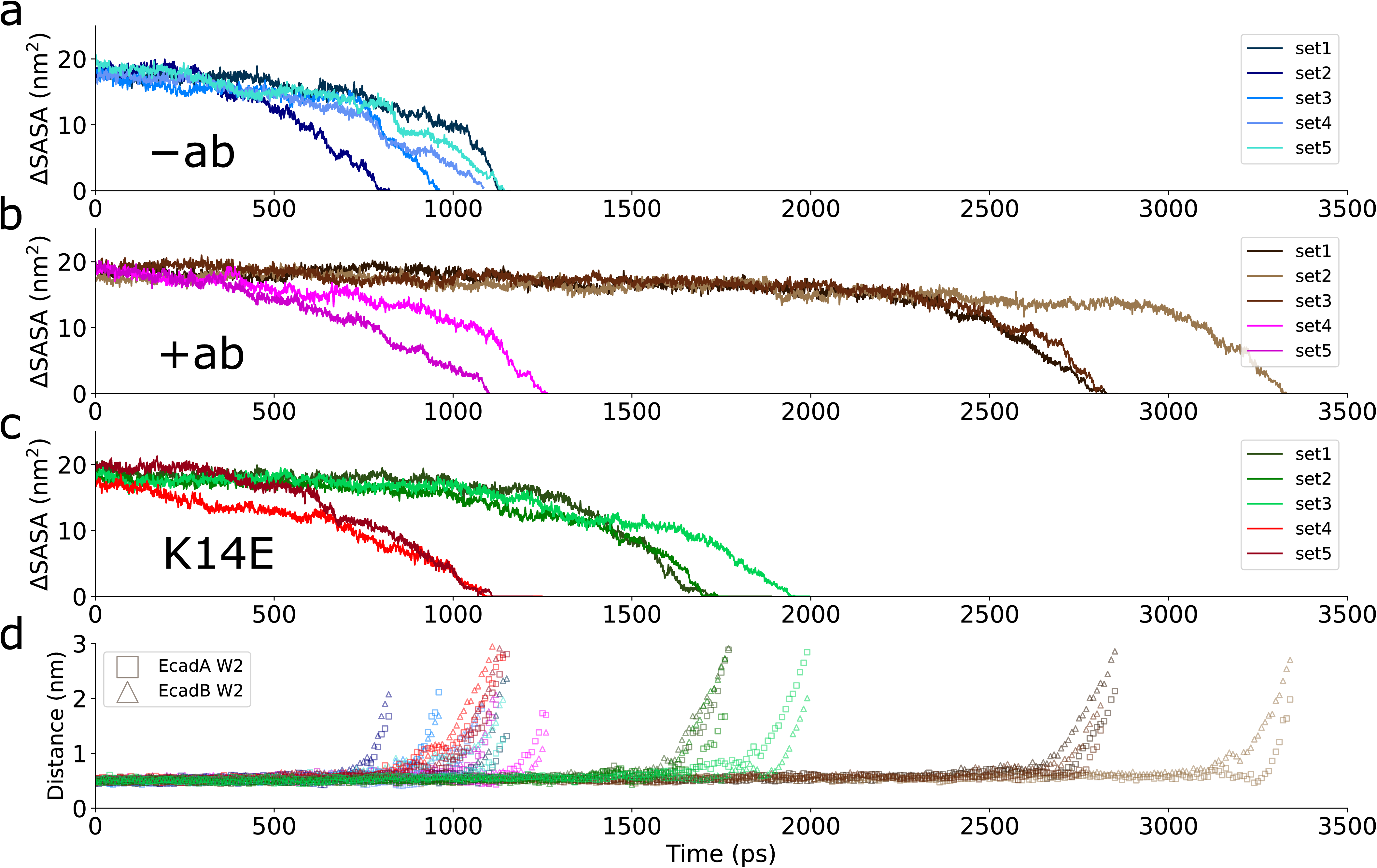
Adhesion strengthening requires two bound 66E8 to form IEC1 with partner Ecad. Constant-force SMD simulations with change in Ecad-Ecad interfacial area calculated from ΔSASA in the (a) WT Ecad −ab condition, (b) WT Ecad +ab condition, and (c) K14E +ab condition. While the lifetimes of the Ecad–Ecad bonds are similar in the WT −ab, sets 4-5 of the WT +ab condition, and sets 4-5 of the K14E +ab condition, the lifetime of the Ecad–Ecad bond in sets 1– 3 of the WT +ab condition, where both interacting Ecads form IEC1 with 66E8, are substantially longer. The lifetimes of K14E-K14E interactions are slightly longer when both K14E mutants form IEC1’ with 66E8. (d) Distance between center of mass of W2 and the center of mass of the hydrophobic pockets in each of the constant-force SMD simulations. In every SMD simulation, the time point that the W2 leaves its binding pocket is almost identical to the time that ΔSASA drops to zero, suggesting that the primary barrier for the dissociation of the Ecad strand-swap dimer remains W2’s exit from its binding pocket.

We also performed MD and SMD simulations on Ecad strand-swap dimers bound to a single 66E8 Fab (Supplementary Fig. S2a, “1ab” condition). Consistent with our previous finding that Ecad strand-swap dimers bound to only one activating antibody cannot strengthened binding (Xie et al., 2022), in the 66E8 1ab condition, the ΔSASA also reached zero at ∼800ps, indicating binding cannot be strengthened by only one 66E8 (Supplementary Fig.S2b).

The presence of a point mutation (K14E) in Ecad decreased the binding strengthening effects of 66E8. The SMD simulations showed two almost indistinguishable populations: while sets 1, 2 and 3 of the K14E mutants interacted for ∼1700ps suggesting slightly stronger interactions, the K14E mutants in sets 4 and 5 only remained bound for ∼1100ps (Fig. 3c).

As an additional measure of strand-swap dimer stability, we calculated the distance between the center of mass of W2 and the center of mass of its complementary binding pocket during every constant force SMD simulation (Fig. 3d). These measurements provide insights into the duration of each W2’s retention within the hydrophobic pocket, indicating the persistence of the β-strands in their swapped position. Consistent with the ΔSASA analysis, we observed that in sets 1-3 of the +ab condition, where Ecad strand-swap dimers were stabilized, W2s were retained in its hydrophobic pocket for a longer period of time compared to all other simulation conditions (Fig. 3d). In every SMD simulation, the time point that each W2 leaves its binding pocket was almost identical to the time that ΔSASA dropped to zero, suggesting that 66E8 binding does not alter the fundamental nature of Ecad adhesion: the primary barrier for the dissociation of the Ecad strand-swap dimer remained W2’s exit from its binding pocket.

Collectively, our simulations provide substantial evidence to support of our hypothesis that the presence of 66E8 leads to the stabilization of Ecad through the formation of IEC1, resulting in enhanced binding. The introduction of a K14E mutation disrupts IEC1 formation and results in only a marginal reinforcement of Ecad binding.

### 66E8 inhibits Ecad conformational changes during strand-swap dimer dissociation

The dissociation of strand-swap dimers necessitates a conformational change in Ecad, involving the displacement of W2 from its complementary binding pocket and the "un-swapping" of β-strands. The precise structural transitions involved in the force-induced dissociation of strand-swap dimers and how these conformational changes are influenced by mAbs like 66E8 are unknown. We therefore monitored the conformation of Ecad during SMD simulations, in the presence (+ab) and absence (-ab) of 66E8. Compared to their unstrained state, we observed a distinct flattening of Ecad strand-swap dimers near the onset of dissociation due to the applied pulling force (Fig. 4a for +ab condition and Supplementary Fig. S3a for -ab condition).

**Figure 4.**
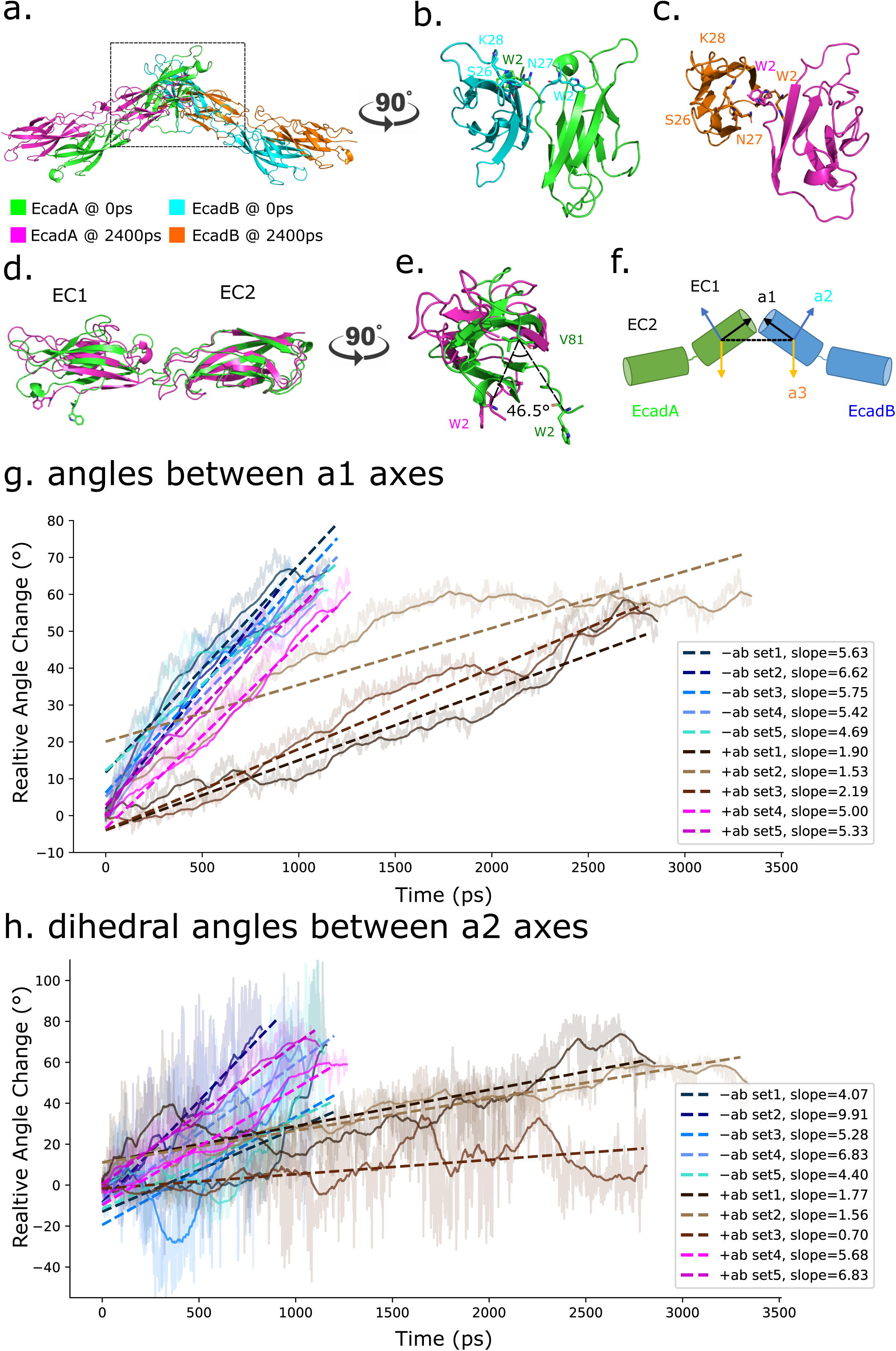
66E8 inhibits Ecad conformational changes which are necessary for the strand-swap dimer dissociation. Force-induced dissociation of Ecad strand-swap dimer requires conformational changes on Ecad EC1 domain which is shown using +ab condition set 1 SMD as an example. For better visibility, only Ecads are shown and 66E8 is not shown. (a) Ecad strand-swap dimers become flattened by force during SMD as shown by the alignment of strand-swap dimer structure at the start of SMD (0ps, EcadA in green, EcadB in cyan) with the structure at the end of SMD (2400ps, EcadA in magenta, EcadB in orange). (b) Side view of the strand-swap dimer shows W2 are blocked by the partner Ecad S26-K28 loop in the absence of force (0ps). (c) Side view of the strand-swap dimer shows EC1s are rotated near the end of SMD simulation (2400ps) so that W2 is no longer blocked by the partner Ecad S26-K28 loop. (d) Alignment of single Ecad at the start of SMD (0ps) and near the end of SMD (2400ps) indicates that while EC2 domains remain almost unchanged, there are conformational changes in EC1 domains. (e) W2s rotate ∼46.5⁰ (calculated with a reference residue V81) during SMD. (f) Scheme for measuring Ecad conformational change during SMD using angle change calculations with EC1s’ principal axes. The flattening conformational change is measured by the angle change between a1 principal axes which are along the longest axes of EC1s (shown as black arrows). The rotation conformational change during SMD is measured by the dihedral angle between a2 principal axes which are perpendicular to the a1 and lie in the plane of the page (shown as blue arrows). The dihedral angle between a2s are calculated around the black dashed line that passes through the EC1s’ center of masses. (g) Relative angle changes between a1 axes and (h) relative dihedral angle changes between a2 axes were measured in every SMD simulations in both −ab and +ab conditions, and were fitted to linear regression models. The slope in the linear regression model represents the rate as shown in the figure legends (unit is 10^-2^ degrees/ps). While all sets in −ab and sets 4 and 5 in +ab conditions have similar slopes, sets 1-3 in +ab conditions where 66E8 forms IEC1 and stabilizes both Ecads have a much smaller slope. This shows that 66E8 induced Ecad stabilization slows down the conformational change that drives strand-swap dimer dissociation.

A closer look at W2’s escape from its complementary pocket revealed that the EC1 domains rotate during dissociation. In the unstrained strand-swap dimer, residues S26, N27, and K28 within the binding pocket act as a cap, which prevents W2 from escaping (Fig. 4b and Supplementary Fig. S3b). However, as the strand-swap dimer dissociates, the EC1 domains rotate relative to each other, which frees W2 from the S26-K28 loop (Fig. 4c and Supplementary Fig. S3c). Measurement of the angle change between the α-carbons of W2 and a reference V81 revealed a rotation of W2 by ∼46.5⁰ in the +ab condition and ∼37.8⁰ in the -ab condition. Taken together, this suggests that the conformational changes occurring during Ecad strand-swap dimer dissociation involve both the flattening of the overall dimer and the rotation of EC1 domains relative to each other. The EC1 conformational changes in the presence and absence of 66E8 are similar. This suggests that 66E8 binding does not induce these conformational changes, but instead, these conformational changes are the pathway along which strand-swap dimers dissociate.

To examine if 66E8 alters the rate at which these conformational changes occur, we measured the relatively angular change between Ecad EC1 principal axes (a1, a2, and a3) during the SMD simulations (Fig. 4f). First, we measured the rate of strand-swap dimer flattening during SMD by calculating the angle between the principal axes a1 which correspond to the longest axis of Ecad EC1 domain. In agreement with the ΔSASA results, our analysis revealed the presence of two distinct populations. In the first population, which consisted of sets 1, 2, and 3 in the +ab condition where Ecad was stabilized by 66E8, the rate of flattening was significantly slower, ∼1.9×10^-2^ degrees/ps (Fig. 4g, brown). In the second population, which comprised all sets in the −ab condition and sets 4 and 5 in the +ab condition where 66E8 was absent or failed to stabilize the strand-swap dimer, the rate of flattening was approximately 3-fold faster, ∼5.5×10^-2^ degrees/ps (Fig. 4g, blue and magenta).

Next, to investigate the rate of rotational movements of EC1 domains within strand-swap dimers during SMD simulations, we calculated dihedral angles between the principal axes a2 across the line joining the centers of mass of the EC1 domains. This angle provided an estimation of EC1 rotation occurring in the strand-swap dimer during each SMD run. Similar to the “flattening” analysis, our “rotation” analysis revealed the presence of two distinct populations. In the first population, which included sets 1, 2, and 3 in the +ab condition, the rate of rotation was significantly slower, approximately ∼1.3×10^-2^ degrees/ps (Fig. 4g, brown). In the second population consisting of all sets in the −ab condition and sets 4 and 5 in the +ab condition, the rate of rotation was ∼5-fold faster, around ∼6.1×10^-2^ degrees/ps (Fig. 4g, blue and magenta).

In addition, when K14E mutation was introduced, 66E8 binding did not slow down the flattening of the strand-swap dimer or the rotation of EC1 domains relative to each other (Supplementary Fig. S4). Taken together, our simulation results demonstrate that in the binding strengthening mode, where the β-strands and their complementary binding pockets are stabilized, 66E8 significantly slows down the Ecad conformational changes which are necessary for strand-swap dimer dissociation.

### Experimental single molecule measurements show that 66E8 strengthens Ecad binding

We used single molecule AFM force measurements to directly test our simulation predictions that 66E8 binds to Ecad in two modes, only one of which strengthens adhesion. In our experiments, we immobilized full-length ectodomains (EC1-5) of human-Ecad, which were biotinylated at their C-terminus, on AFM cantilevers and glass coverslip substrates functionalized with PEG tethers and streptavidin, following previously described protocols (Koirala et al., 2021; Xie et al., 2022). We then measured Ecad-Ecad interactions in the absence (–ab; Fig. 5a, left panel) and presence (+ab; Fig. 5a, right panel) of 66E8 Fab. For the +ab experiment, Ecads on the coverslip and the cantilever were incubated with 66E8 and AFM experiments were performed in the presence of free 66E8 Fab in the measurement buffer.

**Figure 5.**
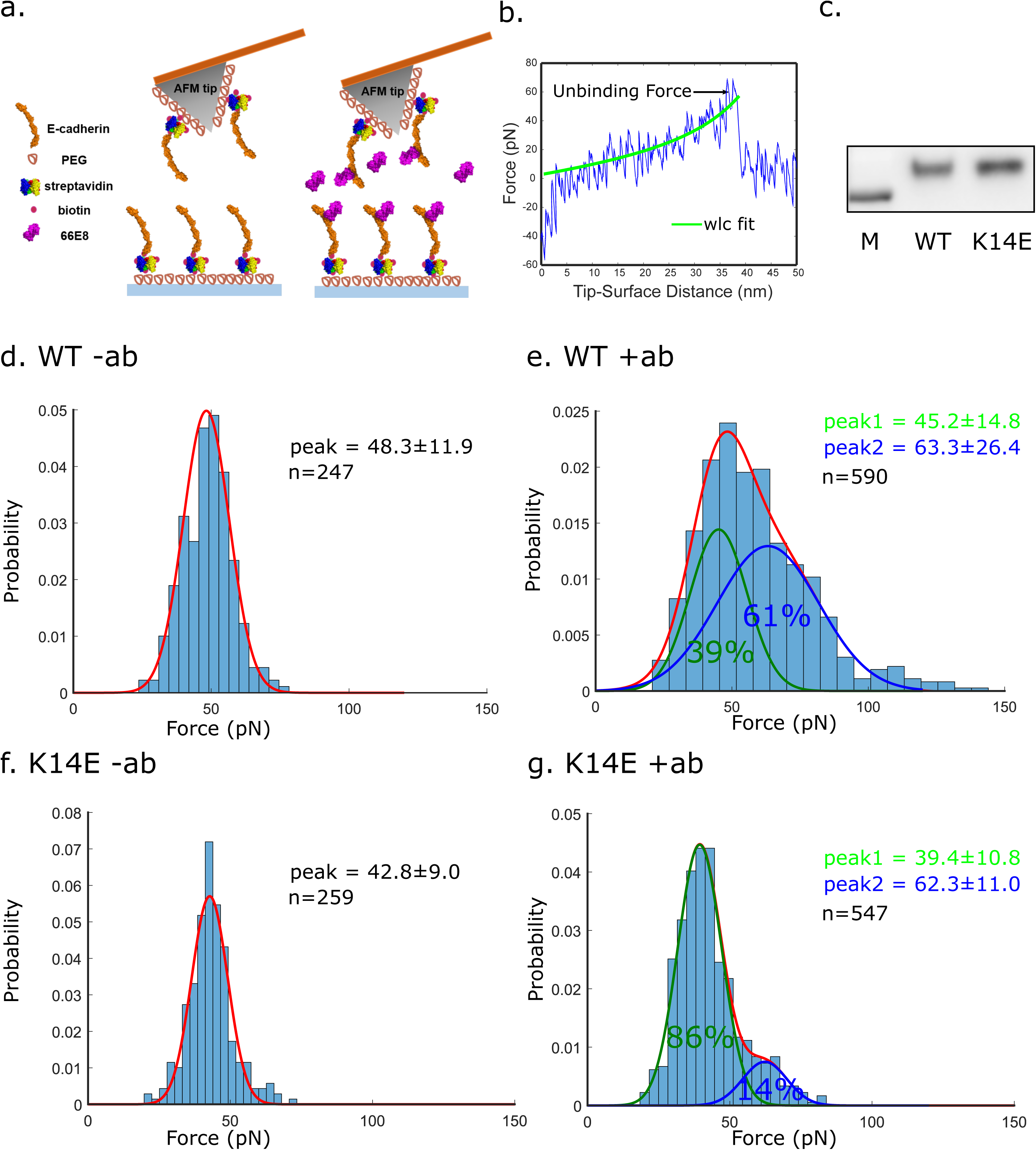
Single-molecule measurements of 66E8-mediated strengthening of Ecad. (a) Left: Scheme for AFM experiments carried out in the absence of 66E8 (−ab). Biotinylated Ecads were immobilized on an AFM cantilever and substrate functionalized with PEG tethers and decorated with Streptavidin. Right: Scheme for AFM experiments carried out with 66E8 (+ab). Both AFM cantilever and substrate were incubated with 66E8 and measurements were carried out with free 66E8 in the buffer. (b) Example force curve. Stretching of the PEG tether, which serves as a “signature” of a single-molecule unbinding event, was fit to a worm like chain (WLC) model (green line). (c) Western blots show that 66E8 recognizes both WT Ecad and Ecad K14E mutants. The molecular weight marker (M) corresponds to a molecular weight of 75 kD. Single molecule AFM force measurements were performed with WT and K14E Ecad in the absence (−ab) and presence (+ab) of 66E8. Histograms of the unbinding forces were generated by binning the data in each condition using the Freedman–Diaconis rule. The optimal number of Gaussian distributions for each fit was determined using BIC. This analysis prescribed one Gaussian distribution for (d and f) and two Gaussian distributions for (e and g). (d) Probability density of Ecad–Ecad unbinding forces measured in the absence of 66E8. Forces are Gaussian distributed (red line) with a peak force of 48.3 ± 11.9 pN. (e) Probability density of Ecad–Ecad unbinding forces in the presence of 40 nM 66E8 was best fit by a bimodal Gaussian distribution. While the first peak at 45.2 ± 14.8 pN (green line) corresponds to un-strengthened Ecad unbinding, a second peak at 63.3 ± 26.4 pN (blue line), which accounts for 61% of the unbinding forces, corresponds to strengthened adhesion. (g) Probability density of K14E–K14E unbinding forces measured in the absence of 66E8. Forces are Gaussian distributed (red line) with a peak force of 42.8 ± 9.0pN. (h) Probability density of Ecad K14E–K14E unbinding forces measured in the presence of 40 nM 66E8 was best fit by a bimodal Gaussian distribution. While the first peak at 39.4 ± 10.8 pN (green line), which accounts for 86% of the unbinding forces, corresponds to un-strengthened K14E-K14E bonds, the second peak at 62.3 ± 11.0 pN (blue line), which accounts for only 14% of the measured unbinding forces, represents strengthened interactions. This indicates that the K14E mutation on the Ecad IEC1 interface, diminishes the strengthening effects of 66E8.

In our experiments the AFM tip and substrate were brought into contact to allow the Ecads to interact homophilically. Subsequently, the cantilever was withdrawn from the substrate at a constant velocity (1 μm/s), and the Ecad-Ecad unbinding force was recorded. Specific single molecule unbinding events were identified from the non-linear stretching of PEG tethers; the PEG stretching data was fit to the worm-like chain (WLC) model using least square method (Fig. 5b). Prior to the AFM experiments, we confirmed that 66E8 recognized the Wild Type (WT) and K14E mutant Ecads using western blots (Fig. 5C). The unbinding forces histograms were fitted to Gaussian distributions; the optimal number of Gaussian distributions for each experimental condition was predicted using the Bayesian information criterion (BIC, Supplementary Figure S5).

In the absence of 66E8 (–ab), the unbinding force histograms were best described by a single Gaussian distribution with a peak unbinding force of 48.3 ± 11.9 pN (Fig. 5d). However, when the experiments were performed in the presence of 40nM 66E8, the unbinding force distributions exhibited a bimodal behavior. While the lower force peak corresponded to the un-strengthened – ab condition (45.2 ± 14.8 pN; Fig. 5e), the second peak had a higher unbinding force corresponding to strengthened Ecad-Ecad bonds (63.3 ± 26.4 pN; Fig. 5e). As an additional experimental test, we performed the AFM experiments at a different pulling velocity (5 µm/s, Supplementary Fig. S6). At a higher pulling velocity, we again observed a unimodal force distribution in the –ab condition and a bimodal gaussian distribution in the +ab condition.

To test the predicted role of K14 on 66E8 mediated adhesion strengthening, we generated human Ecad K14E mutants. We used single molecule AFM force measurements to test the adhesion of the human Ecad K14E mutants in the presence (K14E +ab) and absence (K14E –ab) of 66E8. Since the K14 residue in Ecad is not located within the 66E8 binding site, the K14E mutation did not have any effect on the ability of 66E8 to recognize Ecad, as confirmed by western blot analysis (Fig. 5c). In the absence of 66E8, the unbinding forces between K14E-K14E bonds were best described by a single Gaussian distribution, resulting in an average unbinding force of 42.8 ± 9.0 pN (Fig. 5f). While we observed a bimodal force distribution when the experiments were performed in the presence of 40 nM 66E8 Fab, the unbinding force histogram was dominated by a large peak at a lower force (Fig. 5g). The lower force peak, which accounted for 86% of the unbinding forces, corresponded to un-strengthened K14E-K14E bonds at 39.4 ± 10.8 pN. The second peak, which accounted for only 14% of the measured unbinding forces, represented strengthened interactions at 62.3 ± 11.0 pN (Fig. 5g). These findings align with our simulation predictions, demonstrating that Ecad K14E mutants significantly diminish the strengthening effect of 66E8, reducing the population with enhanced binding from 61% to only 14%. Furthermore, our AFM results validate the existence of the IEC1 since even though the Ecad K14 residue is not part of the 66E8 binding site in the crystal structure, the K14E mutation still impacts the adhesion strengthening function of 66E8.

## Discussion

Using MD simulations, SMD simulations, and AFM experiments, we have demonstrated that the formation of a novel IEC1 interface between 66E8 and the Ecad EC1 domain plays a crucial role in stabilizing both the swapped β-strand and the pocket region. Consequently, in order to enhance Ecad adhesion, the IEC1 interface must be established between the strand-swap dimer and 66E8, on both Ecads. Destabilizing this interface, as observed with the K14E mutant, leads to a reduction in binding reinforcement. Due to the stochastic nature of IEC1 formation, 66E8 exhibits interactions with Ecad in two distinct modes: one that strengthens the bond between Ecad and another that does not alter Ecad binding.

Importantly, our findings provide a molecular rationale for why antibody mediated stabilization of the swapped β-strand and the hydrophobic pocket region, strengthens Ecad binding. Our data shows that the activating antibodies significantly impede Ecad conformational changes, which are necessary to rupture the strand-swap dimer. Dissociation of the strand-swap dimer occurs when each W2 residue escapes from its binding pocket, causing the dimer to flatten and the Ecad EC1 domains to rotate relative to each other. However, 66E8 stabilizes the conformation of the strand-swap dimer and restricts conformational changes, thereby strengthening Ecad binding. It is possible that the cryo-EM structure showing a twisted conformation of the full-length Ecad ectodomain bound to 19A11 captures a similarly strengthened strand-swap dimer conformation (Maker et al., 2022).

Additionally, while previous studies identified key residues in the W2 hydrophobic pockets which affect the binding affinity of cadherins (Vendome et al., 2014a), we show that a loop formed by amino acids S26-K28, serves as a cap which prevents W2s from disengaging from their complimentary binding pockets. When Ecad EC1 domains rotate with respect to each other, the loop S26-K28 no longer blocks W2 from escaping.

It is noteworthy that the binding interface observed in the x-ray crystal structure does not reveal the mechanism by which 66E8 strengthens Ecad adhesion. Previous studies show that crystal structures only reflect a snapshot of protein-ligand complexes at cryogenic temperatures, where the absence of thermal energy and pulling forces prevent the complex from adopting their functional conformation at physiological temperatures (Zheng et al., 2015). Consequently, there are many examples of crystal structures that fail to capture physiologically relevant receptor-ligand conformations (Davis et al., 2008). Fortunately, our atomistic simulations successfully predicted the formation of an interface between 66E8 and the Ecad EC1 domain, which was not observed in the crystal structure. Most importantly, we experimentally validated these predictions using site directed mutagenesis and single molecule AFM force measurements.

Besides AFM experiments, we also performed cell-based adhesion activating experiments using Colo 205 cells expressing WT human Ecad or K14E Ecad mutants (Supplementary Fig.S7). Colo 205 cells are routinely used to test the function of adhesion activating mAbs such as 19A11 and 66E8, which induce rounded Colo 205 cells to adopt an epithelial morphology (Petrova et al., 2012). By adding 66E8 Fab to the cell media, we showed that 66E8 stimulates Ecad mediated adhesion and triggers compact epithelial morphology in Colo 205 cells expressing WT human Ecad. In contrast, 66E8 did not activate adhesion in Colo 205 cells expressing Ecad K14E mutants (Supplementary Fig.S7). While this result can be interpreted to show that the K14E mutation diminishes 66E8 mediated adhesion strengthening, it could also arise because the K14E mutation abrogates adhesion in cells (Harrison et al., 2010).

In addition to forming robust strand-swap dimers, Ecads also dimerize in a weaker X-dimer conformation. X-dimers, which are mediated by a salt bridge between K14 and D138, are thought to serve as intermediates in the pathway leading to the formation of strand-swap dimers (Harrison et al., 2010; Li et al., 2013; Manibog et al., 2016; Sivasankar et al., 2009) and subsequent dissociation on the cell surface (Hong et al., 2011). Unlike 19A11 which directly binds to K14 and inhibits the formation of X-dimers, 66E8 binding does not block access to the K14 residue, as evidenced by our western blot data showing that 66E8 still recognizes the Ecad K14E mutant. However, the crystal structure of the 66E8-Ecad complex suggests that the 66E8 sterically hinders the formation of X-dimers (Maker et al., 2022). Nonetheless, Ecad bound to 66E8 still adopt a strand-swap dimer conformation either because 66E8 rearranges its position to allow transient formation of X-dimers or because 66E8 binds to Ecad only after strand-swap dimerization is complete.

It is also possible that blocking X-dimer formation could prevent strand-swap dimer dissociation and thereby strengthen adhesion. Our data shows that since a key interaction in IEC1 involves K14, formation of IEC1 may also inhibit dissociation of Ecad by preventing X-dimerization. While our study does not directly investigate the role of X-dimers during Ecad unbinding, our data suggests that the force-induced dissociation of strand-swap dimers do not involve an X-dimer intermediate. Our AFM experiments demonstrate that the K14E mutant, where X-dimer formation is blocked, has similar unbinding force as Wild-Type Ecad (Figs. 5d and 5f) and our SMD simulations show that Ecad strand-swap dimers do not adopt an X-dimer conformation during dissociation. Finally, in addition to binding in *trans* conformations, Ecads on the same cell surface also form *cis* dimers (Harrison et al., 2011). However, 66E8 binding does not block the *cis*-dimer interface and consequently is unlikely to interfere with *cis*-dimer formation.

While our study shows that 66E8 strengthens Ecad ectodomain adhesion independent of the cytoplasmic region, previous studies show that Ecad adhesion on the cell surface can be regulated by intracellular proteins such vinculin (Koirala et al., 2021), and p120-catenin (Petrova et al., 2012; Shashikanth et al., 2015). Previous single molecule studies show that vinculin regulates Ecad adhesion via changes in actomyosin contractile forces, which switch Ecad ectodomains from X-dimer to strand-swap dimer structures (Koirala et al., 2021). Similarly, the phosphorylation state of p120-catenin has been shown to regulate Ecad mediated cell adhesion, and Ecad activating antibodies have been shown to induce p120-catenin dephosphorylation (Mendonsa et al., 2020; Petrova et al., 2012). It is possible that p120-catenin dephosphorylation also stabilizes the strand-swap dimer by modulating intracellular contractile forces, although the mechanism by which this occurs is still unknown.

Our study demonstrates that 66E8 strengthens Ecad ectodomain adhesion using similar biophysical mechanisms as the adhesion activating mAb 19A11. Both 66E8 and 19A11 form electrostatic interactions with the K14 residue on the Ecad EC1 domain and selectively stabilize the swapped β-strand and the pocket region. This stabilizes the strand-swap dimer which consequently impedes conformational changes that are necessary for Ecad dissociation. Our results demonstrate that selectively targeting the structural determinants of strand-swap dimer formation are sufficient to strengthen cadherin adhesion. We anticipate that these insights will serve as guidelines for design of new mAbs to enhance Ecad adhesion.

## Methods and Material

### Generation of WT Ecad, K14E Ecad, and 66E8 Fab for AFM experiments

The ectodomains of human wild-type Ecad (residues 1-709) were cloned with Avi-tag and 6XHis tag at the C-terminal and incorporated into pcDNA3.1(+) vector as previously described (Xie et al., 2022). Single point mutation K14E was introduced using NEB Q5 site-directed mutagenesis kit. The full plasmids were expressed in HEK293T cells through transient transfection with PEI (Milipore Sigma). After 3-5 days post transfection, conditioned media with added protease inhibitor (Thermofisher Scientifics) was collected, and stored at 4°C. His tagged WT and K14E Ecad were affinity purified using a gravitational column containing Ni-NTA agarose beads (Qiagen). The Ni-NTA beads were then washed with biotinylation buffer (pH 7.5, 25 mM Hepes, 5 mM NaCl, and 1 mM CaCl_2_) and bound Ecad was biotinylated at 30°C for 1hr using the BirA Enzyme kit (BirA 500 Kit, Avidity). Biotinylated Ecad was eluted with elution buffer (pH 7.5, 500 mM NaCl, 1 mM Ca Cl_2_, 20 mM HEPES, 200 mM Imidazole). Proteins were then dialysed into storage buffer (pH 7.5, 10mM Tris HCl, 100mM NaCl, 10mM KCl, 2.5mM CaCl_2_), flash frozen (ethanol and dry ice), and stored at -80 degrees.

Generation of the 66E8 hybridoma has been described previously (Petrova et al., 2012). 66E8 Fabs were recombinantly generated by cloning the variable regions of the heavy and light chain into the mouse IgG1 Fab backbone (GenScript).

### Western blot

The purified human WT E-cad and K14E E-cad mutant ectodomains were boiled for 5 mins in SDS sample buffer (Bio-rad, 90% 4X Laemmli Sample Buffer + 10% 2-Mercaptoethanol) and run on a 4-15% polyacrylamide gels (Mini-PROTEAN® TGX™ Precast Protein Gels, Bio-Rad) at 200V for 30 mins. Proteins were transferred to a PVDF membrane (Immuno-Blot®, Bio-Rad) at 200 mA for 1 hr and incubated with 5% blocking buffer (PBS + 0.1% tween 20 + 5% blotting-grade blocker) for 1 hr. After washing, the membrane was incubated with 66E8 (1:1000 dilution with blocking buffer) for 1 hr at room temperature followed by incubation with goat anti-mouse IgG (Invitrogen) with HRP conjugate (1:5000 dilution with blocking buffer) for 30 mins at room temperature. WesternBright ECL HRP substrate (Advansta) was used to detect the protein.

### Single-molecule AFM experiments

The biotinylated WT Ecad or K14E mutant ectodomains were immobilized onto the surface of AFM cantilevers (Hydra 2R-50N, AppNano) and glass coverslips (CS) using previous protocols (Koirala et al., 2021; Xie et al., 2022). The cantilever and CS were treated with Piranha solution (25% H_2_O_2_, 75% H_2_SO_4_) overnight and washed with deionized (DI) water. The CS was further cleaned with 1 mM KOH and washed with DI water. The cantilever and CS were then washed with acetone and silanized with 2% 3-aminopropyltriethoxysilane (Millpore Sigma) in acetone solution for 30 mins. After silanization, 10% biotin-PEG-Succiniminidyl (MW5000, Laysan) and 90% mPEG-Succinimidyl Valerate (MW5000, Laysan) was dissolved in 100 mM NaHCO_3_ and 600 mM K_2_SO_4_ to formulate 100 mg/ml PEG incubation buffer. The cantilever and CS were incubated in PEG buffer for at least 4 hrs and washed with DI water.

Prior to start of the experiment, the PEGylated cantilever and CS was first blocked with BSA at 10 mg/ml for 1 hr at room temperature. The cantilever and CS were then incubated with 0.1 mg/ml streptavidin (Thermofisher) for 30 mins and 200 nM Ecad proteins solution for 90 mins. To block the extra biotin binding sites on streptavidin, the cantilever and CS were incubated with 0.02 mg/ml free biotin solution.

All AFM measurements were performed on Agilent 5500 AFM system with a close loop scanner. Force measurements were performed in a buffer (pH 7.5, 10 mM Tris⋅HCl, 100 mM NaCl, 10 mM KCl, and 2.5 mM CaCl_2_) and the spring constant of the cantilever was calculated based on thermal fluctuation method (Hutter and Bechhoefer, 1993). To identify specific unbinding events, all PEG stretching events were fitted to a WLC using least-square fitting protocol. The unbinding events were filtered based on root mean square error of the WLC fitting, and the unbinding forces were measured. The unbinding forces were binned using Freedman-Diaconius rule and fitted using a Gaussian mixture model. The optimal number of Gaussians were determined using Bayesian Information Criterion (BIC).

### MD simulation and analysis

MD simulations were conducted on the FARM high-performance computing cluster at University of California, Davis with GROMACS 2022.3 as previously described (Xie et al., 2022). The simulations were performed in OPLS-AA/L force field (Kaminski et al., 2001) and TIP4P water model with 10Å radius cut-off for Van der Waals and electrostatic interactions. To calculate the electrostatic energy, we used the particle mesh Ewald method with a 0.16-grid spacing. At the beginning of the simulation, the Ecad crystal structure (PDB 2O72) or the Ecad crystal structure bound to 66E8 (PDB 6VEL) was placed in the center of a dodecahedral box so that every atom was at least 1 nm from the boundary and the system was relaxed with energy minimization and stabilized with equilibration under isothermal-isochoric and isothermal-isobaric conditions using a modified Berendsen thermostat and Berendsen barostat. The box was filled with water molecules and neutralized with charged ions (100 mM NaCl, 4 mM KCl, and 2 mM CaCl_2_). After stabilization, 60 ns MD simulation was performed with 2-fs integration steps. The protein structure usually reached equilibration after ∼20 ns. The C-α RMSF of each residue in the Ecad EC1 domain (residues 1–100) during the final 35 ns MD was calculated using the *gmx rmsf* module. The distances between charged atoms for the hydrogen bonds K105:H110 and distance between interface residues’ center of masses were calculated using the *gmx pairdist* module.

### Constant-force SMD simulations and analysis

The constant force SMD simulation was performed on the FARM high-performance computing cluster as described previously (Xie et al., 2022). The starting structure for SMD simulation was the last frame of the corresponding MD simulation. This structure was placed in the center of a rectangular box such that the longest axis of the structure was parallel to the longest axis of the box and no atom was closer than 1 nm to the boundary (30 × 12 × 8 nm for the −ab conditions; 30 × 15 × 15 nm for the +ab/1ab conditions). The system, containing ∼38000 atoms for −ab condition and ∼88000 atoms for +ab condition, was relaxed and equilibrated under isothermal-isobaric conditions using the same protocol as in MD simulation. During each SMD simulation, we fixed the C-terminus of one Ecad and pulled on the last residue on the C-terminus of the other Ecad along the longest axis of the box with a constant force ∼665 pN (400 kJ⋅mol−1⋅nm−1). The solvent assessable surface area (SASA) was calculated using *gmx sasa* module, and the changes in solvent accessible surface (ΔSASA) was calculated using the equation: ΔSASA = SASA [protein A] + SASA [protein B] − SASA [protein A + protein B]. The Ecad EC1 principal axes were obtained using *gmx principal* module. Both angles between a1 axes and dihedral angles between a2 axes were calculated using the equation: *angles* = 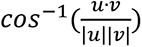, where u, v represent the normalized vectors of principal axis.

## Acknowledgments

Research supported by the National Institute of General Medical Sciences of the National Institutes of Health (R01GM133880 to SS and R35GM122467 to BMG).

## Author Contributions

B.X., S.S., and B.M.G. designed research. B.X., and S.X. performed research. B.X., and S.X. analyzed data. B.X., S.X. and S.S. wrote the paper.

