## Supplemental Information for "Strengthening E-cadherin adhesion via antibody mediated binding stabilization"

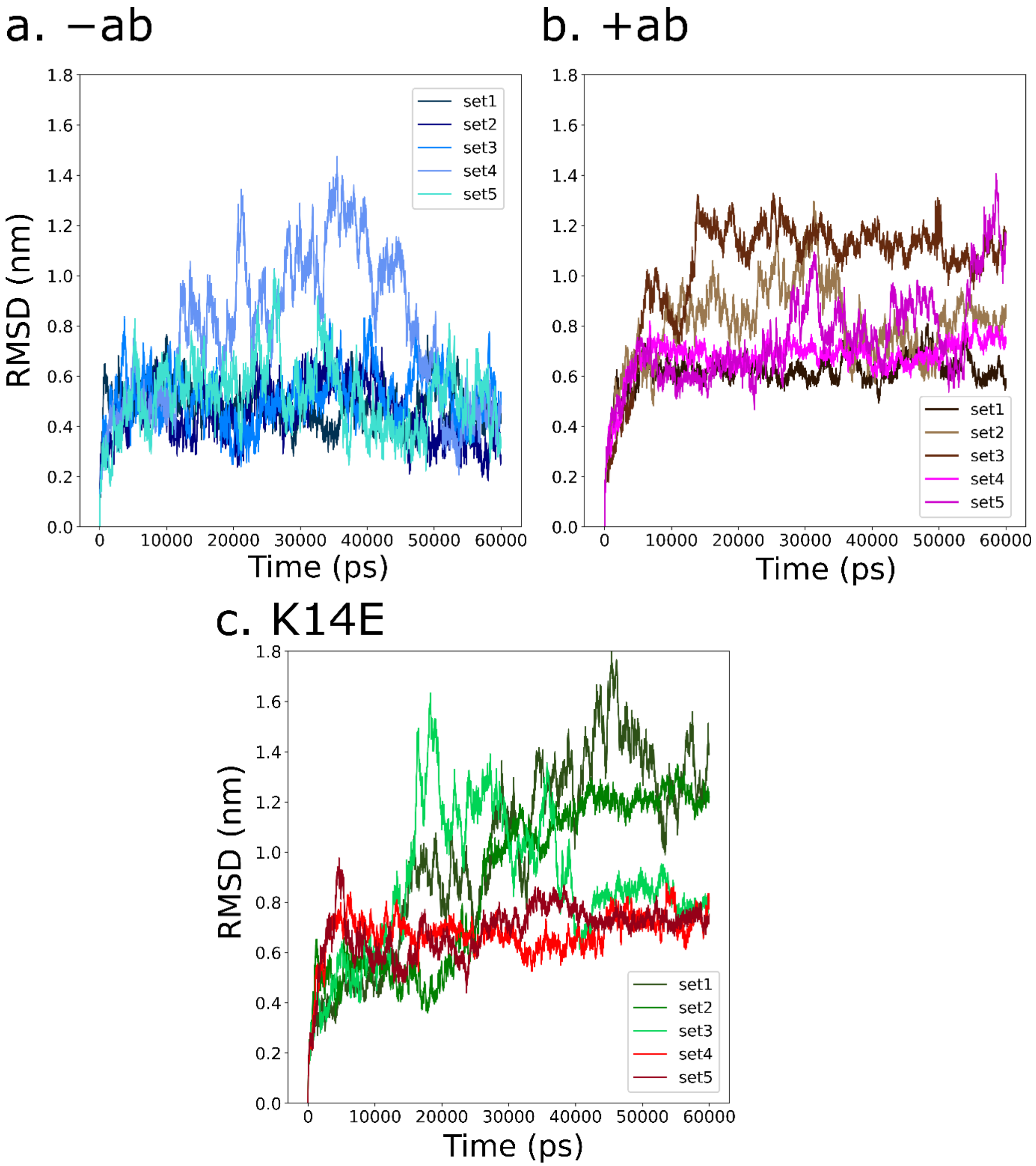


**Fig. S1. Protein backbone RMSD in MD simulations relative to the initial structures at the start of simulation.** RMSD values measured for each set in three conditions (a) −ab condition, (b) +ab condition, and (c) K14E +ab condition. Stabilization of RMSD values suggest that structures are well equilibrated.

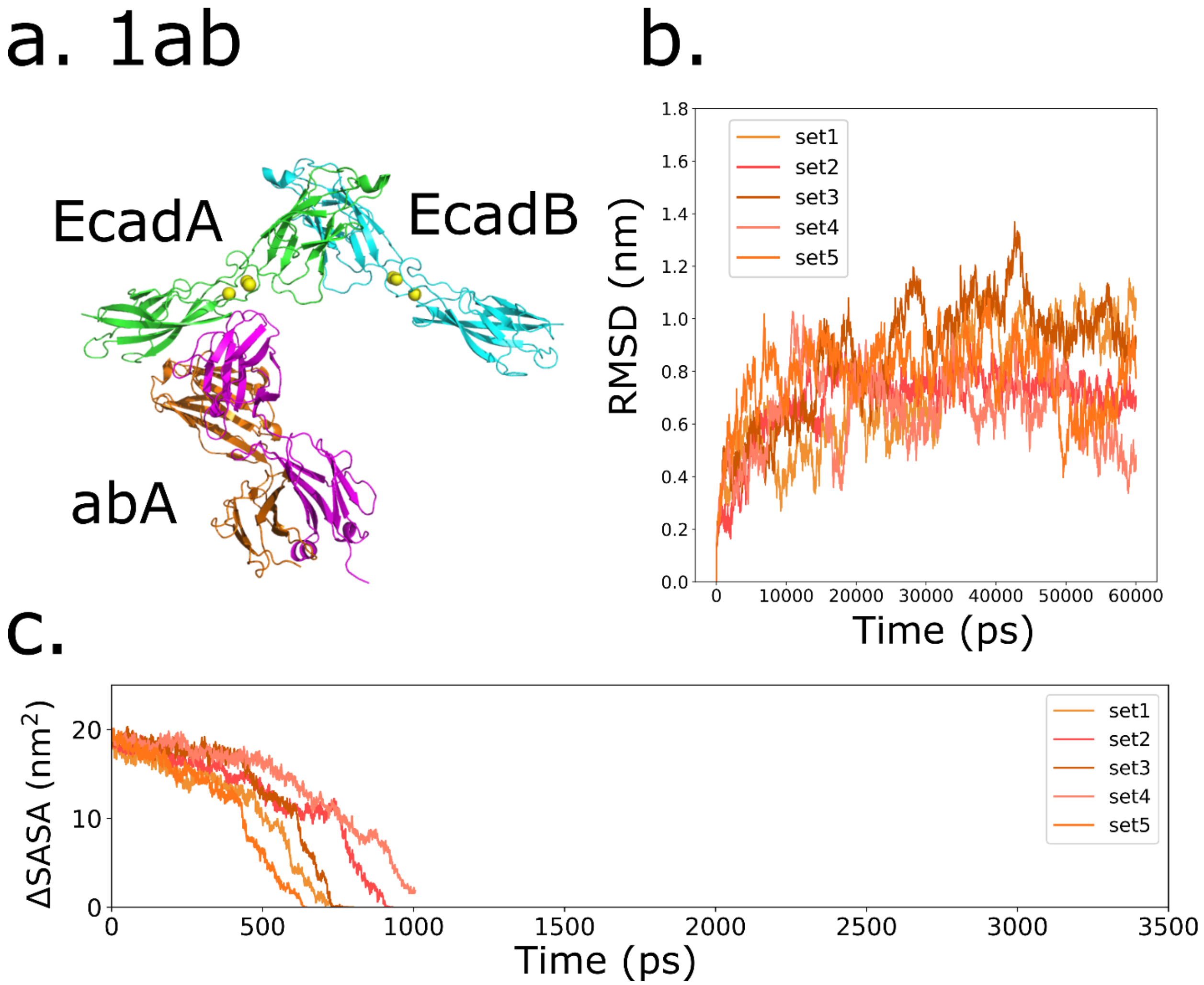


**Fig. S2. Ecad strand-swap dimers cannot be strengthened by single 66E8 binding.** (a) Ecad strand-swap dimer bound to a single 66E8 (abA) on one of the Ecad (EcadA). This 1ab structure was used as the starting structure in MD and SMD simulations. (b) Protein backbone RMSD of every 1ab MD simulation relative to the initial structure at the start of simulation. Stabilization of RMSD values suggest that structures are well equilibrated. (c) Constant-force SMD simulations of 1ab condition, with change in Ecad-Ecad interfacial area calculated from ΔSASA. Ecad-Ecad interactions lasted ~800ps, which is similar to the Ecad interactions without 66E8 (−ab condition), suggesting that a single 66E8 cannot strengthen Ecad strand-swap dimers.

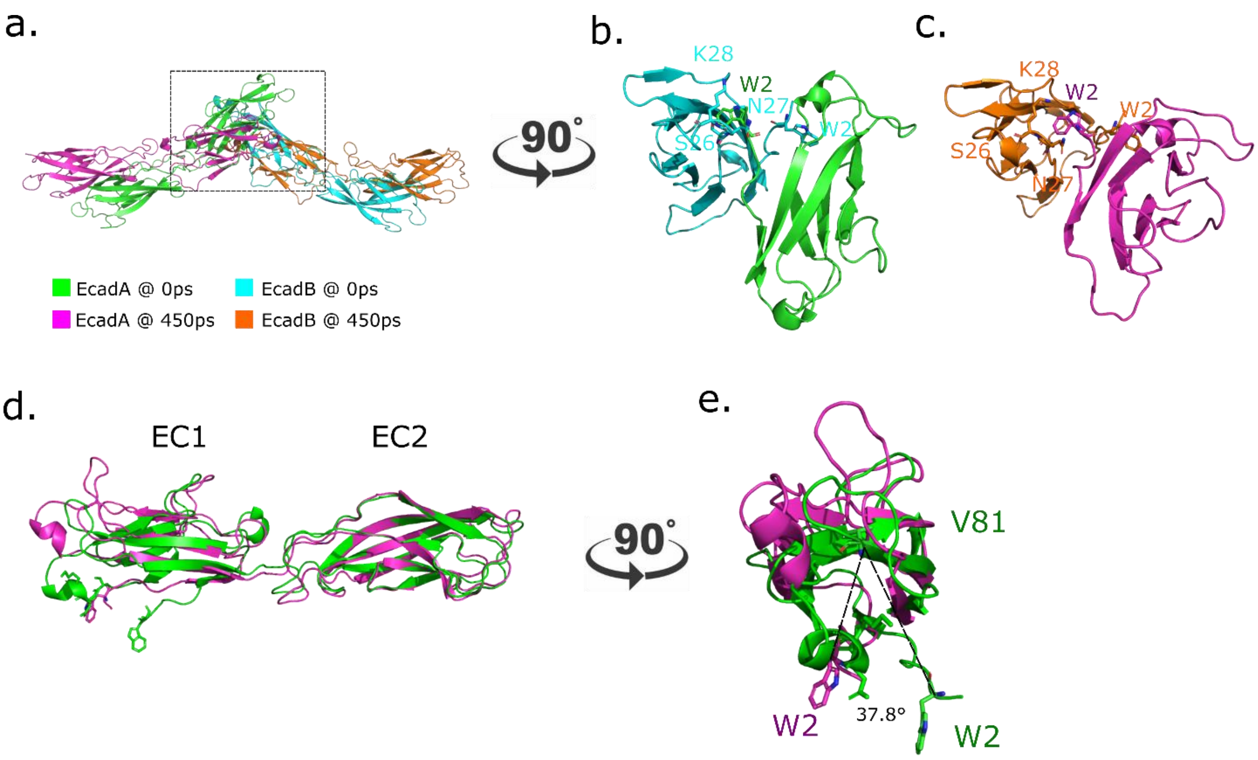


**Fig. S3. Ecad strand-swap dimer conformational changes during force-induced dissociation.** The force-induced dissociation of Ecad strand-swap dimer requires conformational changes on Ecad EC1 domain which are shown using set 1 in −ab conditions SMD as an example. (a) Ecad strand-swap dimers become flattened by force during SMD as seen from the alignment of strand-swap dimer structure at the start of SMD (0ps, EcadA colored in green, EcadB colored in cyan) with the structure at the end of SMD (450ps, EcadA colored in magenta, EcadB colored in orange). (b) Side view of the strand-swap dimer shows that escape of W2s are blocked by the partner Ecad S26-K28 loop in the absence of force (0ps), while (c) side view of the strand-swap dimer shows EC1s are rotated so that escape of W2s are no longer blocked by the partner Ecad S26-K28 loop near the end of SMD (450ps). This suggests that a rotation of EC1 domains is necessary during dissociation. (d) The alignment of single Ecad at the start of SMD (0ps) and near the end of SMD (450ps) indicates that while the EC2 domain remains almost unchanged, there are conformational changes in the EC1 domain. (e) The W2 residue rotates ~37.8º (calculated with a reference residue V81) during SMD.

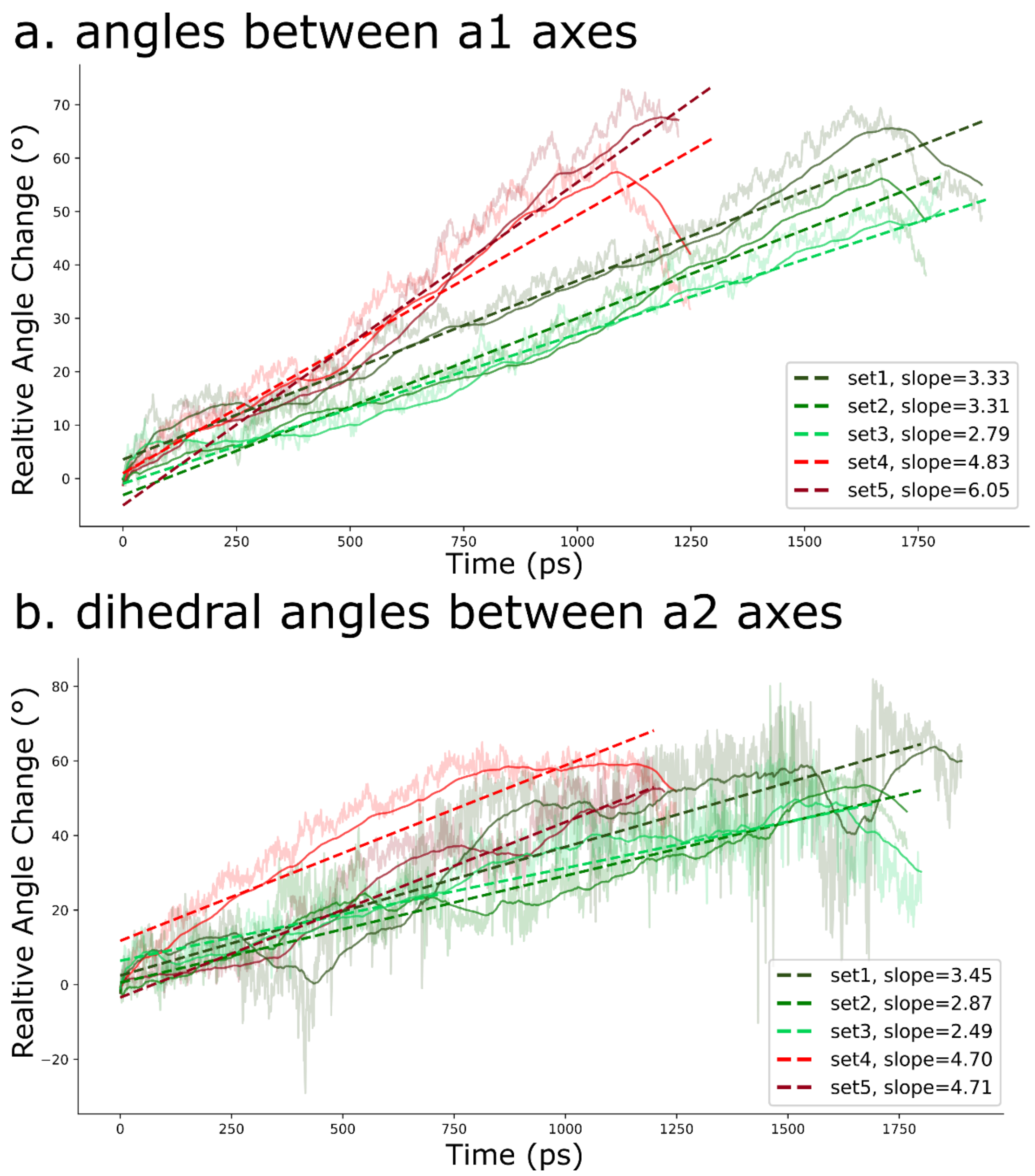


**Fig. S4. Characterizing Ecad K14E mutant conformational changes during SMD.** (a) Relative angle changes between the a1 axes and (b) relative dihedral angle changes between a2 axes measured in every SMD simulations in “K14E + ab” condition. Data was fit to a linear regression model. The slope in the linear regression model represents the changing rate are shown in the figure legends (unit is $10^{-2}$ degrees/ps).

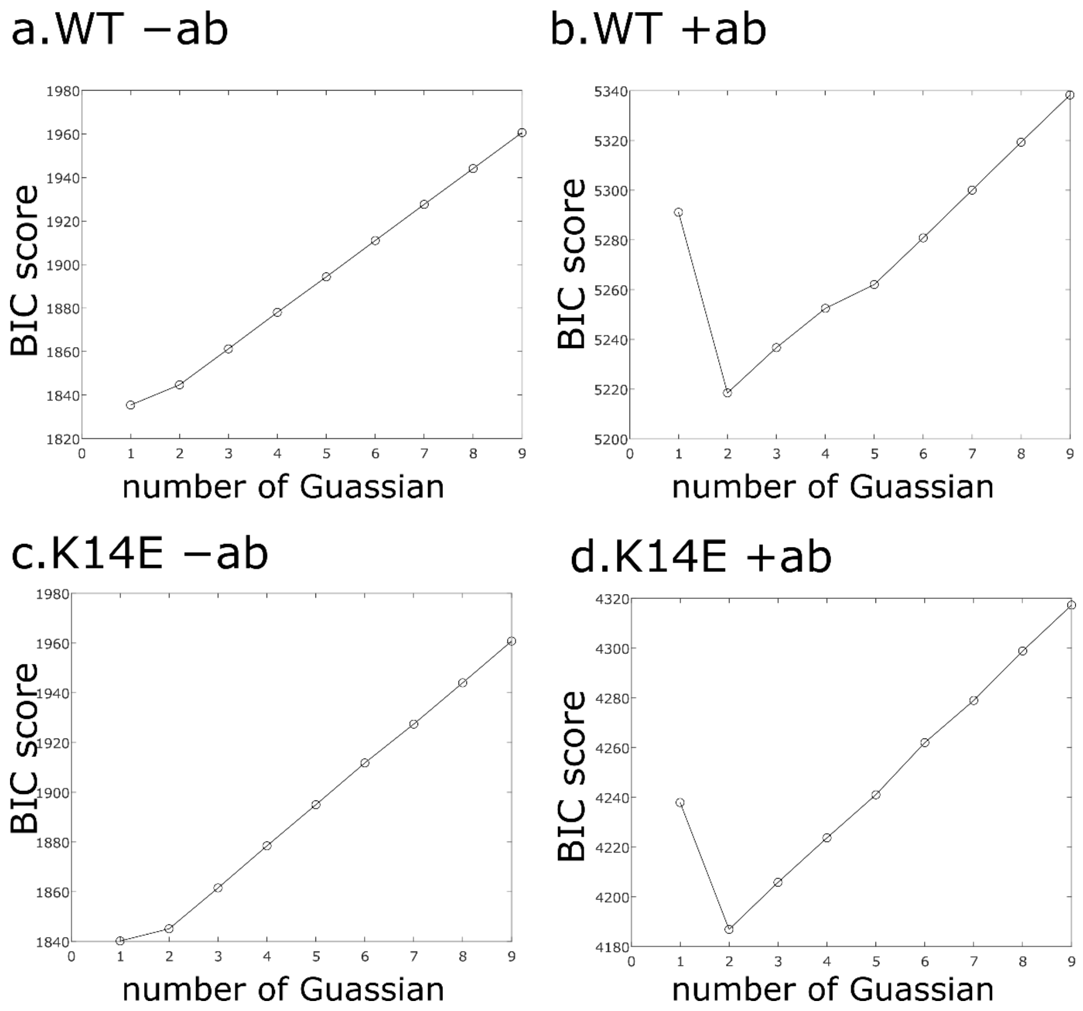


**Fig. S5. Bayesian information criterion (BIC) predicts the optimal number of Gaussians in the unbinding AFM force histograms.** BIC ascribes a penalty to reflect the fact that more free parameters will always yield a better fit; lower BIC scores correspond to a more likely model. A single Gaussian distribution optimally describes (a) WT-Ecad and (c) K14E Ecad. A bimodal Gaussian distribution best describes (b) WT +ab and (d) K14E +ab, force distribution.

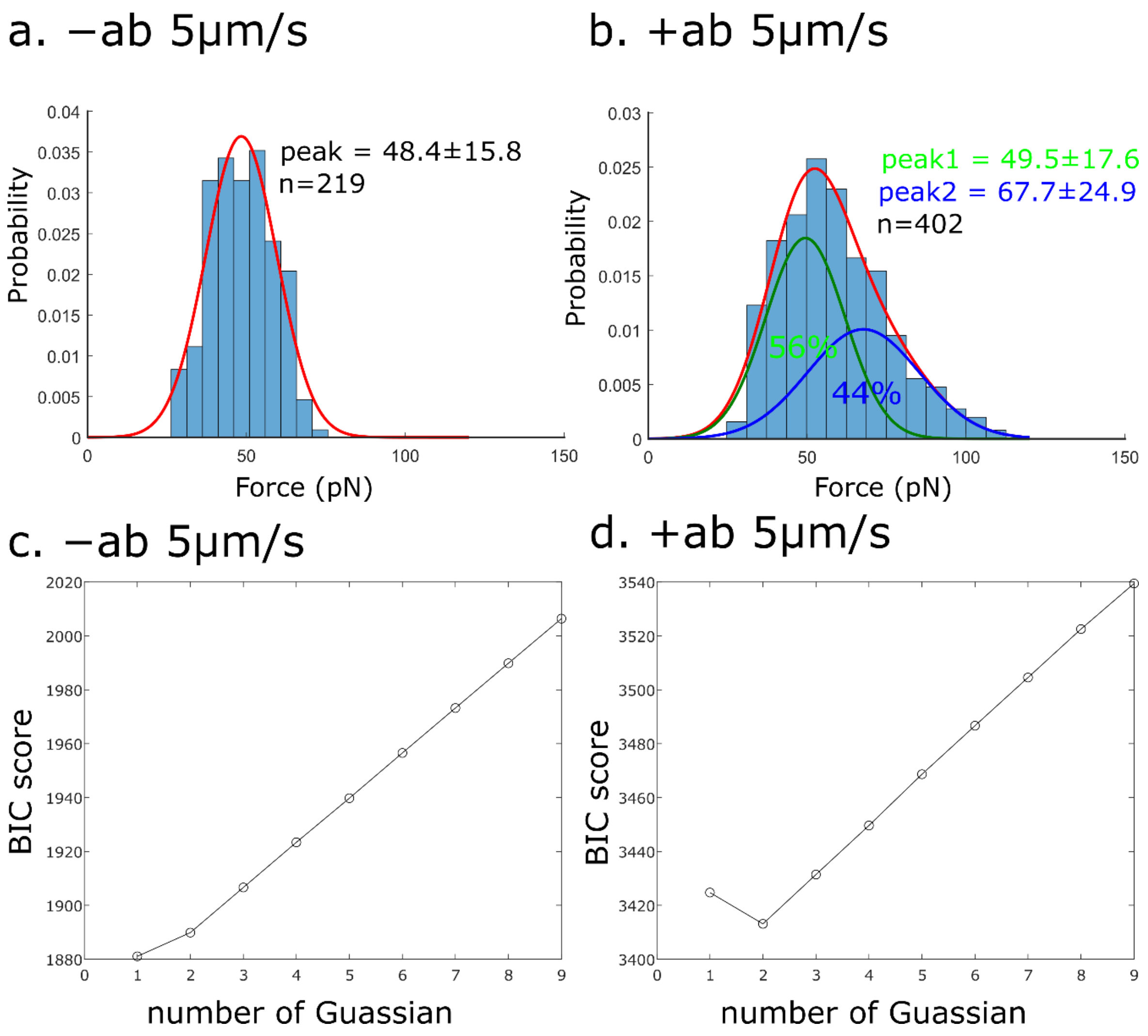


**Fig. S6. Atomic force microscopy (AFM) experiments performed using 5μm/s pulling velocity.** Experiments were performed using wild-type (WT) human Ecad in the (a) absence ("−ab") and (b) presence ("+ab") of 66E8. Histograms of the unbinding forces were generated by binning the data in each condition using the Freedman–Diaconis rule. The optimal number of Gaussian distributions for each fit was determined using BIC which are shown in (c) and (d). Lower BIC score indicates a more likely model. A single Gaussian distribution optimally describe WT Ecad interactions in the absence of 66E8 with peak force 48.4±15.8pN. A bimodal Gaussian distribution described WT Ecad interactions with 66E8. While the first peak at 49.5 ± 17.6 pN (green line) corresponds to a "native" Ecad unbinding force, a second peak at 67.7 ± 24.9 pN (blue line) corresponds to ~44% the unbinding events strengthened by 66E8.

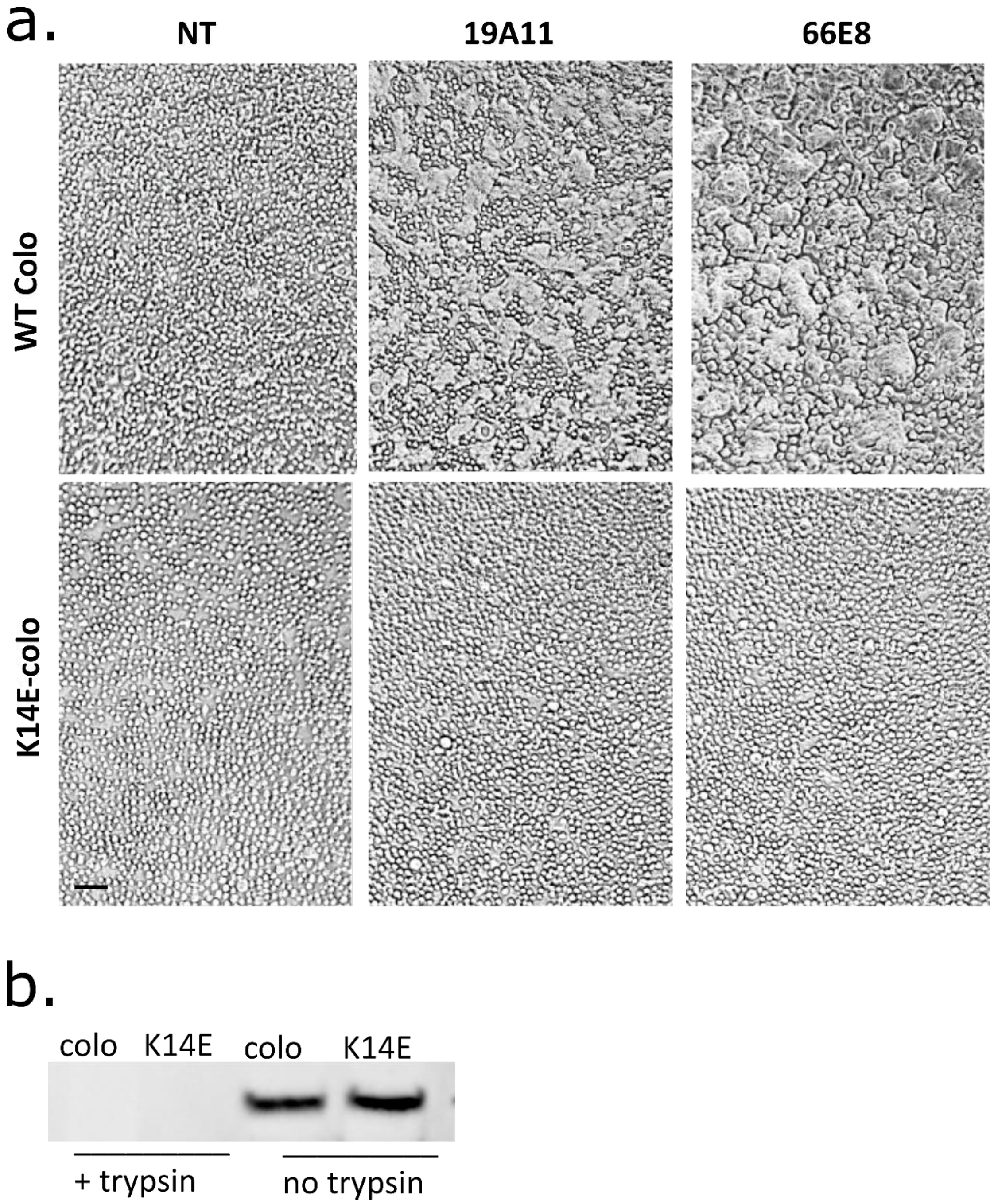


**Fig. S7. 66E8 activates Colo 205 cells expressing WT Ecad but does not activate Colo 205 cells expressing Ecad K14E mutants.** (a) Compared to untreated cells (left column), activating mAbs 19A11 (center column) and 66E8 (right column) triggered compact epithelial morphology in Colo 205 cells expressing WT Ecad (top row). In contrast, 19A11 and 66E8 do not activate adhesion in Colo 205 cells express Ecad K14E mutants (bottom row). Scale bar: 50 µm. (b) Western blots show that WT Ecad or K14E mutants are expressed on the cell surface in Colo 205 cells. Treatment with trypsin digests these cell surface proteins.

### Supplementary movie

**Movie S1. Example constant-force SMD simulations**. One constant-force SMD example from each condition: −ab (set 1), +ab weak conformation (set 5), +ab strong conformation (set 1), and +ab K14E conformation (set 1) are shown. Ecad strand-swap dimers in the −ab, +ab weak conformation, and +ab K14E conformation break significantly faster compared with the +ab strong conformation. Color scheme: Ecad (cyan and green), 66E8 heavy chain (orange), and 19A11 light chain (magenta).

### Supplementary method

**Colo 205 Activation Assay:** The Colo205 activation assay was performed as described previously (Petrova et al., 2012). Briefly, Colo205 cells were densely plated on 96-well plates precoated with 0.1µg/mL rat-tail collagen (Sigma-Aldrich) overnight and treated with activating concentrations of Fabs for 5 hours. Activation was determined by the extent of a morphological change from round cells with distinct borders to a compact epithelial appearance and loss of obvious cell borders.
